# Milpa polyculture enhances productivity through crop complementarity and potential carry over effects

**DOI:** 10.64898/2026.08.17.745153

**Authors:** Carlos Bustos-Segura, Patrick Grof-Tisza, Camilo Rivera, Karlijn de Groot, Raúl González-Salas, Ted CJ Turlings, Betty Benrey

## Abstract

Polycultures have long been practiced in traditional agriculture, yet their ecology-based benefits have remained underexplored. Here, under realistic conditions, we experimentally evaluated the productivity and ecological interactions in cultivated milpa, a traditional Mesoamerican polyculture of maize, squash and beans, using a substitutive design in which total plant density was held constant while varying species composition. Specifically, we asked whether productivity gains arose through complementary or selection effects, and whether these gains were associated with changes in arthropod communities and herbivory. We additionally evaluated whether prior cultivation influenced maize performance in the following season. Milpa plots produced significantly higher total yields, more than 2.6 times those of monocultures, despite poor bean performance. In particular, squash and maize equivalent yields increased approximately threefold. We found that these improvements were mainly explained by complementary effects rather than selection effects. Arthropod communities responded in species-specific ways to crop diversity, with predator abundance tracking herbivore presence. However, no consistent patterns emerged between herbivore load, predator abundance and plant damage, suggesting that belowground plant interactions may play a more important role than top-down herbivore control in explaining complementarity effects. In the following season, maize yield increased by ∼30% in plots previously planted with squash or beans, with milpa plots showing intermediate responses. These findings demonstrate that milpa can substantially enhance productivity while generating benefits that extend into the advantages and soil into the following growing season. Overall, our results suggest that complementarity among crops is the primary driver of productivity in milpa under low-input conditions.

## Introduction

Polycultures, or crop-mixing, are agricultural practices that combine two or more crop species in the same space and time. While many polycultures have deep roots in traditional farming systems, others are modern innovations designed to enhance sustainability. Increasingly, polycultures are promoted as alternatives to industrialized monocultures which produce negative impacts on the environment, with demonstrated benefits for productivity (Iverson et al. 2014), biodiversity (Martinez et al. 2024) and ecosystem functioning (Haddad et al. 2011; Chu et al. 2017). Despite these ecological benefits, evidence that diversification consistently increases crop yield remains mixed and often depends on how productivity is measured (Beillouin et al. 2021).

Traditional polycultures provide some of the clearest examples of how crop diversity can enhance ecosystem functioning. One such system is the milpa polyculture, most commonly composed of maize, beans, and squash. Developed in Mesoamerica thousands of years ago, the milpa integrates three of the region’s earliest domesticated crops and has long been recognized for its resistance to pests and pathogens, reduced dependence on synthetic inputs, and ability to maintain soil quality while providing a nutritionally balanced diet at relatively low cost (Zizumbo-Villarreal et al., 2012; Fonteyne et al., 2023; Vazeux-Blumental et al., 2024; Benrey et al., 2024). Moreover, milpa systems remain widely practiced by smallholder farmers in North and Central America, reflecting their continued ecological and agronomic relevance in low-input agricultural systems.

Polycultures can increase crop productivity through two main mechanisms. While complementarity effects (CE), where interactions among crop species increase community-level productivity, are frequently reported in intercropping systems (Li et al., 2023), selection effects (SE), where plants with higher productivity dominate the output, can also be relevant (Loreau and Hector 2001). Complementarity is commonly attributed to resource partitioning among crops, such as differences in rooting depth or canopy structure that allow plants to exploit distinct ecological niches (Zhang et al. 2014; Yang et al. 2022; Wu et al. 2023). In addition, complementarity in crop mixtures may also arise from ecological interactions that extend beyond direct plant resource use. However, the ecological mechanisms underlying these effects are often inferred rather than explicitly tested. Diversified cropping systems often support greater abundance and diversity of arthropods, including natural enemies of herbivores, which can reduce herbivore pressure through top-down control (Trenbath 1993; Letourneau et al. 2009). Despite these well-documented ecological responses, relatively few studies have simultaneously evaluated arthropod communities and crop productivity under field conditions. For example, a meta-analysis by Letourneau et al. (2009) evaluated whether increased natural enemy diversity associated with producer-level diversity suppresses herbivores but did not assess corresponding effects on crop yield.

Several studies have provided robust evidence of productivity in the milpa from controlled field experiments (Altieri et al. 2012; Zhang et al. 2014; Ebel et al. 2017; Cryan et al. 2025). It has been suggested that differences in root architecture and function among milpa crops can enhance nutrient uptake and contribute to belowground complementarity (Postma and Lynch 2012; Zhang et al. 2014). In addition, recent work has shown that the milpa alters arthropod communities and plant-insect interactions (Liao et al. 2024; Grof-Tisza et al. 2024a). However, previous work focused primarily on insect communities and plant defence traits, including our own (Grof-Tisza et al. 2024a), and did not evaluate plant growth or productivity. As a result, the relative contribution of aboveground ecological interactions and crop complementarity to productivity remains unclear.

Another potentially important but underexplored factor that can contribute to the milpa benefits are soil legacy or carry over effects, by which plants induce changes in soil properties, such as microbiota, chemistry and structure, that persist over time and affect the performance of plants during the following season (de Vries et al. 2012; Ristok et al. 2023). While crop rotation is a well-established practice to mitigate nutrient depletion and enhance productivity (Zhao et al. 2020; Janssen et al. 2026), there is growing evidence that crop mixtures can also influence the soil properties across seasons (Xiao et al. 2022). For example, legumes can contribute to nitrogen availability in ensuing seasons, through symbiosis with rhizobia (Homulle et al. 2022). Because the milpa includes crops with highly distinct belowground traits (Martínez-Romero 2003; Gfeller et al. 2023), it raises the question of whether such complementarity could generate carry over effects in this polyculture system.

Here, we performed a field experiment in native agroecosystem for the three milpa crops in southern Mexico (Fig. 1), using a substitutive experimental design in which plant density was held constant. We tested whether crop diversity in the milpa system generates productivity advantages relative to monocultures while evaluating ecological processes that may contribute to these effects. We quantified productivity using the land equivalent ratio (LER) and net effect ratio (NER), which was further partitioned into complementarity and selection effects. We also characterized arthropod native communities and quantified herbivore damage to evaluate whether changes in insect community structure associated with the milpa polyculture contribute to complementarity effects. Finally, we conducted a follow-up experiment to test whether previous crop combinations influence the performance of a subsequent maize crop potentially through cultivation-dependent carry over effects. By measuring productivity, arthropod communities, and responses in the following growing season, we were able to evaluate multiple mechanisms proposed to explain the benefits of milpa cultivation.

Based on previous work in diversified cropping systems, we expected polycultures to exhibit overyielding relative to monocultures, driven largely by complementarity effects. We also expected crop mixtures to support fewer herbivores and greater abundance of natural enemies. If ecological interactions contribute to such complementarity effects, changes in insect community structure should coincide with differences in crop performance. In addition, if plant–soil conditioning influences crop growth, soils previously associated with crop mixtures should enhance the performance of subsequent plants relative to soils conditioned by monocultures.

**Figure 1.**
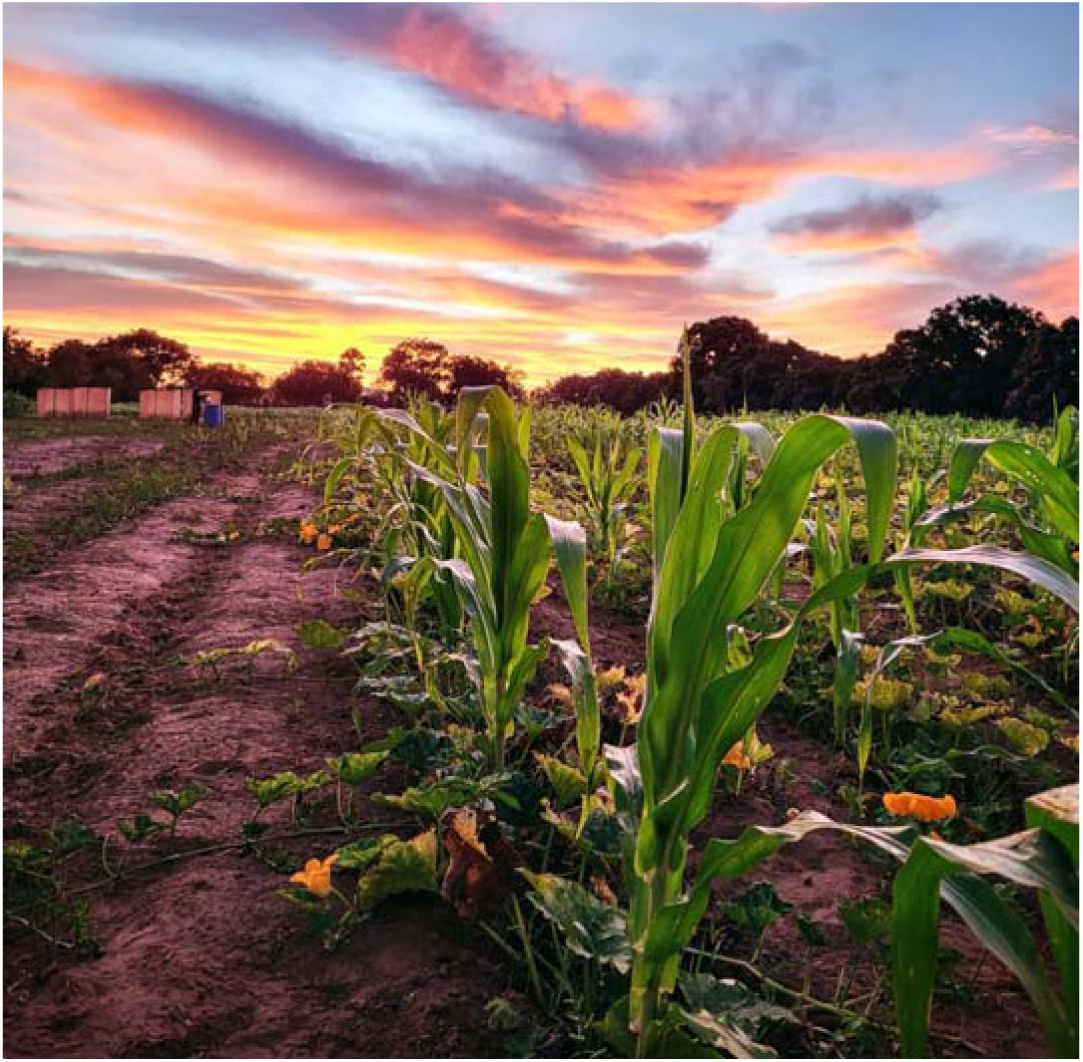
Picture of the experimental field in southern Mexico showing a milpa plot and the rest of the field in the background. (Photograph by Patrick Grof-Tisza).

## Material and methods

### Experimental design

The experimental field was located in an agricultural area surrounded by dry tropical forest in southern Mexico close to the coastal town of Puerto Escondido, Oaxaca. We created plots consisting of monocultures of maize, beans or squash and milpa plots with a mix of the three crops. The experimental field (42 m by 90 m) was divided in four blocks (21 m by 45 m) to account for spatial variability. Each block consisted of 18 plots (7 m by 7.5 m), totalling 72 plots, with seven rows of 7.5 m length per plot. Within each block we planted six plots with milpa while four plots were designated for each monoculture of beans, maize and squash. Cropping treatments were randomly assigned within a block. Planting started at the beginning of November 2022 (start of the dry season). In the previously tilled field, three seeds were manually sown per plant spot. Plants were spaced 1 m apart between rows and 30 cm within rows. We used a substitutive design in which total plant density was kept constant between monocultures and milpa plots, while varying the proportion of crop species. This design allows for comparisons between monocultures and polycultures without confounding diversity effects with differences in overall plant density (Iverson et al. 2014). In milpa plots, crop species were alternated within rows, ensuring heterospecific neighbours on both sides of each plant. One week after germination, seedlings were randomly thinned to leave a single plant per planting spot. Watering was provided through an irrigation system every other day.

### Arthropod community

We conducted two surveys of the arthropod community associated with the plants, one and two months after germination (early December 2022 and early January 2023). These time points were selected as representative measures of the community build-up during the vegetative stage and peak abundances during the pre-flowering stage, based on a previous study (Grof-Tisza et al. 2024a). For each survey, we recorded observations on nine plants in one intermediate row per plot. Plants were selected using a predefined sequence of plant position within the row, consistent across all plots and excluding edge plants to minimize border effects.

Based on previous sampling and updated species lists, we classified arthropods into 93 morphospecies across the following orders: Coleoptera (19), Hemiptera (30), Dermaptera (1), Orthoptera (9), Hymenoptera (19), Mantidae (1), Neuroptera (1) and Araneae (13). Each morphospecies was characterized as either an herbivore or predator based on their observed field activity. Observations were conducted by visually inspecting each individual plant for two minutes and recording the number of individuals per morphospecies in direct physical contact with the focal plant. Each experimental block was surveyed by a different experienced observer.

### Plant traits

Plant size and herbivore damage were measured at two time points during plant development, in late November and December 2022, using the same nine plants selected for arthropod surveys. Plant size in beans and maize was measured from the soil surface to the tip of the longest apical meristem. In squash, size was estimated by summing the lengths of all branches. To estimate herbivore damage, the number of leaves per plant was recorded and the damage per leaf was visually estimated. For bean and squash plants the damage was estimated in percentage in increments of 5%. For maize, the damage was estimated as a damage category of 0 to 4 according to the scale suggested elsewhere (Toepfer et al. 2021). Average damage per plant was calculated as the sum of damage across all leaves divided by the total number of leaves. At the end of the season, squash fruits from each plot were harvested, counted and weighed. Mature maize cobs and bean pods were collected from the central five rows of each plot, then counted and weighed.

### Productivity estimates

Plot yield (in kilograms) was calculated for squash directly from harvest data and for maize and beans by averaging yield from the central five rows and scaling it to the full seven-row plot. To compare yields between monocultures and polycultures with reduced planting density per species, we estimated the expected yield in milpa based on monoculture productivity and the relative proportion of each crop species in the mixture. Total yield per plot was calculated as the sum of yields for all plants present within a plot for both monocultures and milpa plots.

To evaluate productivity differences among plant diversity treatments, we calculated the land equivalent ratio (LER), which quantifies the amount of monoculture land required to achieve the same yield as a polyculture under similar conditions. Values of LER > 1 indicate greater productivity in polycultures relative to monocultures (Li et al. 2023). We also calculated the net effect ratio (NER), which compares the observed productivity of crop mixtures with the productivity expected from monocultures after accounting for the relative proportion of each species in the mixture (Cardinale et al. 2007; Li et al. 2020a). Following Loreau & Hector (2001), we further estimated the net effect of diversity (NE) and partitioned it into selection effects (SE) and complementarity effects (CE). Selection effects arise when highly productive species dominate mixtures, whereas complementarity effects occur when interactions among crop species increase community-level productivity beyond expectations based on monoculture performance.

### Carry over effect of previous crops

A follow-up experiment was conducted during the next wet season (June to August, 2023) to evaluate carry over effects associated with the previous year’s crop diversity treatments. The same field was used and, although the soil was tilled (as is practice) and new planting rows were established, the spatial layout of the original plots was maintained to preserve treatment history.

Maize was uniformly planted across all plots, with rows spaced apart by 1 m and plants separated by 30 cm within rows. In early June 2023, three maize seeds were manually sown at each planting position, and seedlings were later thinned to a single plant. Plants were grown under typical field conditions and received 0.5 g of nitrate fertilizer per plant two and eight weeks after germination. Irrigation was provided primarily by natural rainfall and supplemented only during dry periods exceeding one week. During the experiment, a heavy storm caused partial plant mortality. Therefore, the final number of maize plants per plot was recorded and included in subsequent analyses. At maturity, dried maize cobs were harvested from the five central rows of each plot, pooled, counted and weighed to estimate final yield.

### Statistical analyses

The R software (version 4.3.2) was used for all analyses and plots. We used generalized linear mixed models (GLMMs) with the glmmTMB function (glmmTMB package), and linear mixed models (LMMs) with the function lmer (lme4 package). Statistical significance of explanatory factors was tested with type 2 Wald Chi square tests or F tests depending on model structure.

Arthropod community analyses - Total abundance of herbivores and predators per plant was analysed with repeated measures (GLMMs) with a negative binomial error distribution. Monitoring time, plant species and cropping treatment (monoculture or milpa) and all their interactions were included as explanatory variables. Spatial block and plot position (edge or centre of the field) were used as covariates, and spatial plot was included as a random factor. We estimated as well the Shannon’s diversity index H’ for herbivores and predators separately only for plants that had at least one individual recorded. The variation in arthropod diversity was analysed with an equivalent model as for abundance but with a Gaussian error distribution. To compare the composition of the insect community per plot at the second time point (communities were similar among treatments in the first time point), we used a Redundancy Analysis (RDA) with the rda function (vegan package) with cropping type as the explanatory variable.

Plant traits and yield (first season) - Plant height (maize and beans), squash branch length, and leaf damage were analysed with GLMMs, including monitoring time, cropping type and their interaction as explanatory variable. Spatial block was included as a covariate and plot as a random factor. The yield of each crop was analysed by estimating first the expected yield in milpa based on monoculture productivity per plot. Using linear models, we tested if observed yield per species in milpa was different from their expected yield, including spatial block as fixed factor. We also analysed whether the total yield per plot differed between monocultures and milpa plots. To obtain confidence intervals of productivity indexes we used a bootstrap function with 5000 replicates. We resampled plot-level observations for each diversity treatment per species to estimate LER and NER. Net effects were further partitioned into selection and complementarity effects which were later compared using the bootstrapped distributions.

For the analysing the carry over effects of previous crops, maize height over time was analysed with a repeated measures model (LMM) with monitoring time, soil treatment and their interaction as fixed explanatory factors. Spatial block was included as a covariate, and plot and plant ID as random factors. Then, we compared the number of cobs and total yield per plot among treatments with linear models that included the soil treatment, spatial block and the number of maize plants per plot as explanatory variables.

## Results

### Arthropod community

The abundance of herbivores and predators associated with individual plants was significantly explained by a three-way interaction between plant diversity, plant species and time (Table S1). For the first time point, plant diversity had no significant effect on herbivore or predator abundance for either of the three crop species (P>0.3 for all comparisons). At the second time point, herbivore abundance was higher on bean plants in monocultures than in milpa plots (P<0.0001), whereas the opposite pattern was found for maize (P<0.0001). No difference was detected for squash (P=0.40; Fig. 2a). Predator abundance followed a similar pattern; more predators were found on beans and squash in monocultures compared to milpa (both P=0.04), whereas no significant differences was detected for maize plants (P=0.09; Fig. 2b).

The Shannon’s diversity index H’ for herbivore diversity followed a similar pattern to the herbivore abundance (Fig. 2c) with differences between plant treatment depending on the crop species at the second time point (Table S1). Herbivore diversity on beans was lower in milpa plots compared to monocultures (P=0.0002), whereas the opposite was found in maize plants (P<0.0001) and no differences were detected in squash (P=0.29). In contrast, predator diversity was not significantly affected by any of the experimental factors (Fig. 2d, Table S1). A redundancy analysis (RDA) of arthropod community composition at the second time point revealed that plant diversity explained only 3% of the total variance. Although this effect was small, it was statistically significant (X^2^_(1)_=13.04, P=0.0003). The observed differences were largely driven by two insect species: an unidentified hemipteran specialist on squash that was more abundant in monocultures, and the chrysomelid *Diabrotica balteata* which was more strongly associated with milpa plots.

**Figure 2.**
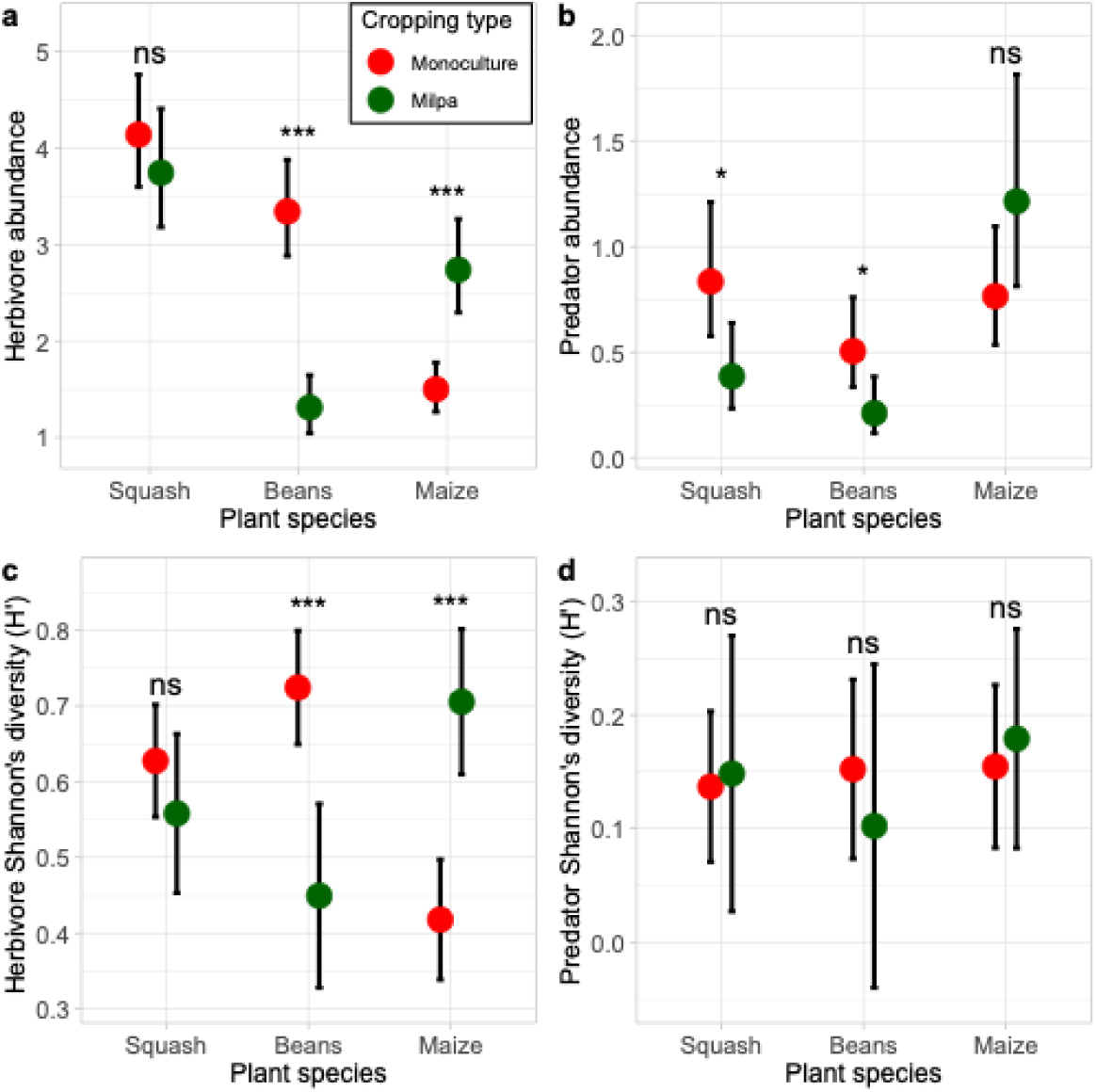
Effect of cropping type on the arthropod community. Herbivore and predator abundance (a and b) and diversity (c and d) associated with individual plants for the three crop species in monocultures and in milpa plots. Panels show estimated marginal means from a GLMM only for the second sampling time; error bars indicate ± 95% C.I. Asterisks indicate the level of statistical significance (* P<0.05, ** P<0.001, ***P<0.0001), ns indicates P>0.05.

### Plant traits

Plant size was not significantly different between diversity treatments for any of the three crop species, but logically plants were larger in the second time point (Table S2). Leaf number increased with time, with no significant effect of plant diversity or squash and maize (Table S2). In beans, however, leaf number not only increased with time, but the interaction between time and diversity treatment was also statistically significant (Table S2). There were no differences in number of leaves between treatments at the first time point (P=0.66), but at the second time point bean plants had significantly fewer leaves in milpa than in monoculture plots (P<0.0001). This reduction in leaf number coincided with high early-season herbivory damage on bean plants. The proportion of leaf area removed by herbivores in squash did not change with time or with diversity treatment (Table S2). In contrast, bean plants experienced greater herbivore damage in milpa plots than in monocultures (Fig. 3b) and this difference was consistent over time (Table S2). For maize, damage levels were similar across treatments but significantly increased over time (Table S2).

### Productivity

Yield responses to crop diversity differed among species (Table S3). Squash and maize showed higher yields in milpa plots than expected from monoculture performance (Figs. 3a, 3b), whereas bean yield was lower in milpa than expected from monoculture performance (Fig. 3c). When the yields of the three species were pooled, total yield per plot in milpa was 2.6-fold higher than monocultures (Table S3; Fig. 3d). Both the land equivalent ratio (LER) (LER=2.14; Table 1) and the net effect ratio (NER=2.88; Table 1) were considerably greater than 1, indicating strong overyielding and a strong productivity advantage of the milpa system.

**Figure 3.**
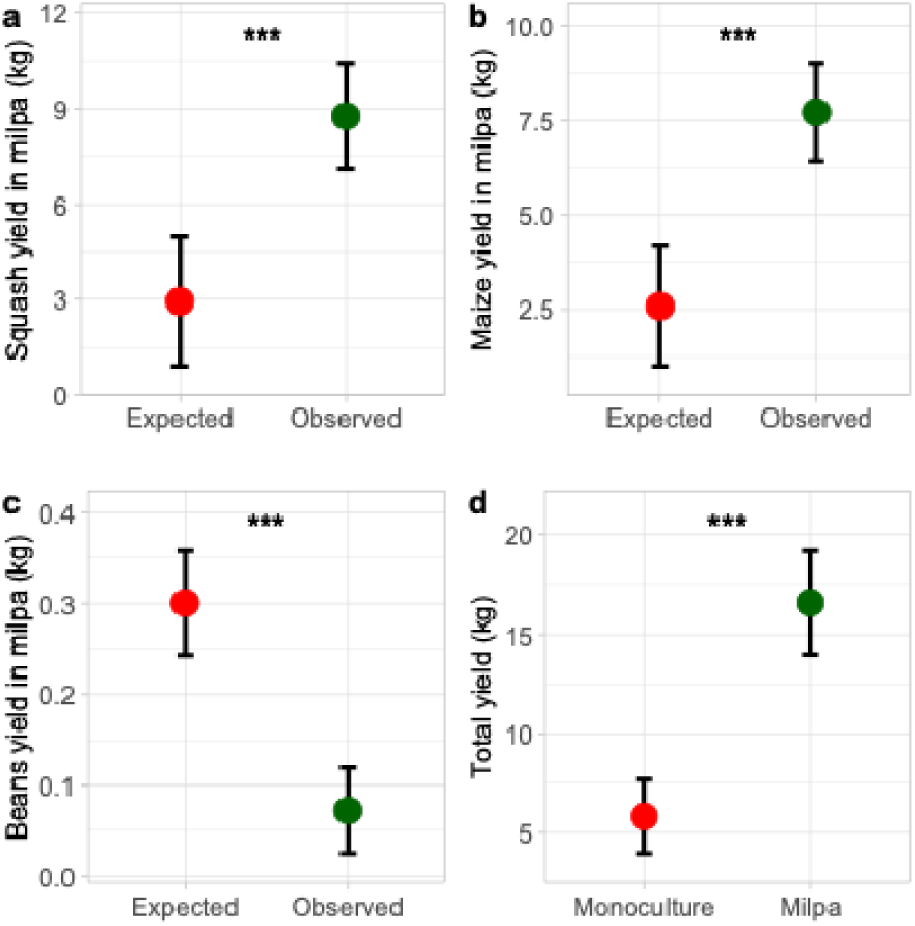
Harvest per plot of each crop species and total yield per plot depending on cropping type. Estimated marginal means of expected and observed yield for squash (a), maize (b), beans (c) in milpa and the total yield of the three crops (d) under monoculture and milpa (polyculture) planting. Expected yield was calculated by adjusting productivity according to the proportion of each species in milpa plots. Error bars indicate ± 95% C.I. Asterisks indicate the level of statistical significance (* P<0.05, ** P<0.001, ***P<0.0001).

We found that the net biodiversity effects (NE) were driven primarily by complementary effects (CE) rather than selection effects (SE), (CE > SE, p = 0.028; Table 1). Accordingly, the proportion of NE explained by CE was higher than the null expectation of 0.5 (mean= 0.6; 95% CI = 0.497, 0.706), indicating that positive interactions among crop species contributed more strongly to productivity than dominance by highly productive species.

**Table 1.** Productivity indices of field experiment comparing polycultures and monoculture plots. Shown are the means and confidence intervals (95% CI) obtained by a bootstrap of 5000 replicates for Land Equivalent Ratio (LER), Net Effect Ratio (NER), Net Effect (NE), Complementarity Effect (CE) and Selection Effect (SE).

| Index | Bootstrap mean | 95% CI |
| --- | --- | --- |
| LER | 2.14 | (1.53, 2.97) |
| NER | 2.88 | (2.06, 3.96) |
| NE (kg/ha) | 2039.81 | (1381.14, 2733.19) |
| CE (kg/ha) | 1231.22 | (701.66, 1797.64) |
| SE (kg/ha) | 808.59 | (560.75, 1031.7) |

### Carry over effects

Maize height was affected by an interaction between time and the crop planted the previous season (Table S4). While pairwise comparisons were not statistically significant, there was a clear trend at 7 and 11 weeks after germination: maize plants grew tallest in plots previously planted with beans or squash, followed by intermediate height in plots previously planted with milpa, and shortest in those with prior maize monoculture (Fig. 4a).

The number of maize cobs was significantly affected by cropping history (Table S4; Fig. 4b). Plots previously planted with squash or beans produced more cobs than those with previous maize; milpa plots had intermediate cob numbers. The cropping type had non-significant effects on maize final yield in the next season (Table S4); however, some trends were observed. Maize tended to be more productive (∼30% higher) in plots where squash and beans were planted the previous season compared to plots previously planted with maize (Fig. 4c), suggesting that crop identity in the previous season can influence maize performance and that the effects of different crop species are largely additive. Maize yield in plots previously planted with milpa was intermediate between the yield in plots previously planted with maize and the other crops.

**Figure 4.**
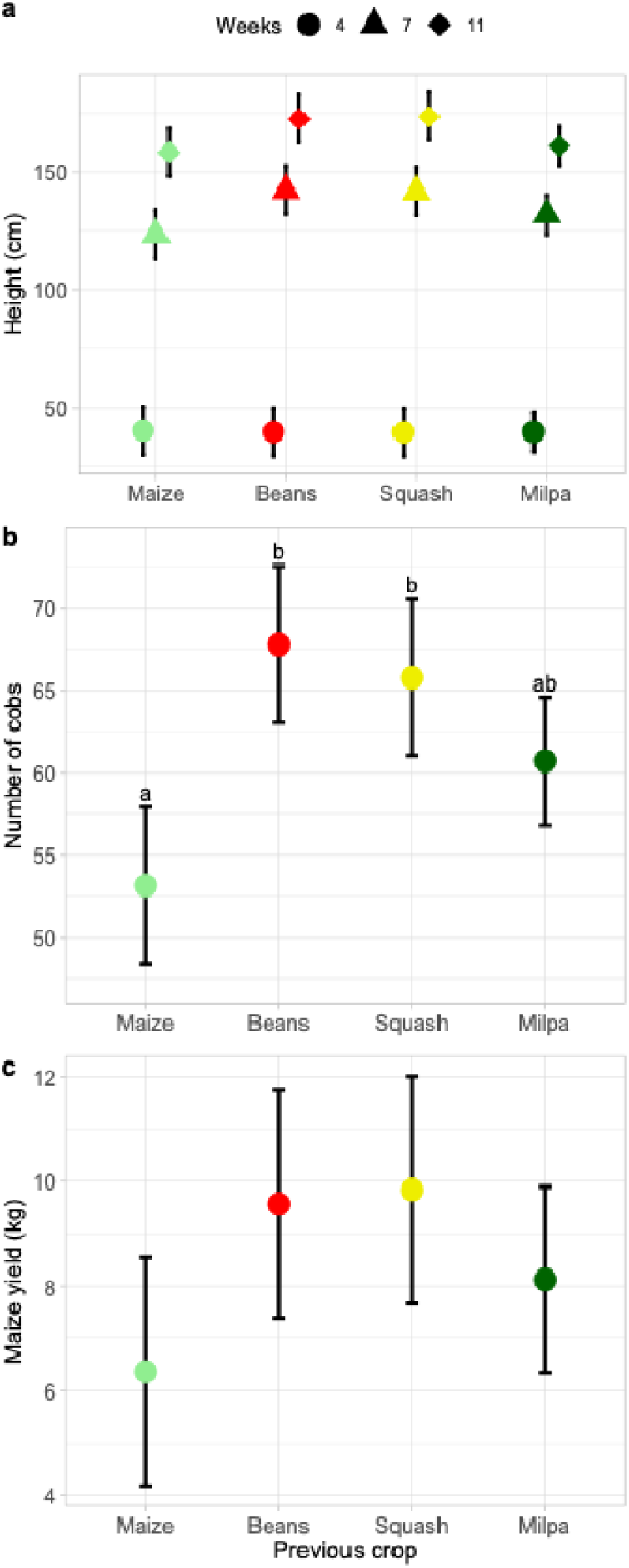
Carry over effects of previous cropping treatments on maize performance. (a) Final maize plant height, (b) number of cobs, and (c) yield per plot in the 2023 wet season. Treatments reflect the crop(s) planted the previous year. Estimated marginal means shown with ± 95% C.I. Different letters between treatment groups indicate significant differences (P<0.05) between pairs of groups.

## Discussion

Agriculture is fundamental for food security, yet current practices continue to drive major environmental challenges, including habitat loss, pesticide overuse, and nutrient pollution. Traditional practices such as crop mixing are receiving renewed attention as potentially more sustainable strategies capable of high productivity while reducing ecological impacts. Despite growing interest in polycultures, relatively few studies have simultaneously evaluated productivity, arthropod communities, and longer-term crop responses under controlled field conditions. Here, using the traditional milpa system as a model, we combined these approaches to evaluate how crop diversity influences crop performance and ecological interactions under low-input conditions.

Using a substitutive design, we compared monocultures and milpa plots while maintaining constant plant density across treatments. Under these standardized conditions, milpa plots more than doubled overall productivity relative to monocultures. Likewise, the LER calculated for the milpa (2.14) was considerably higher than the average reported across intercropping studies (1.23; Li *et al*., 2023), and also higher than previous estimates for the milpa using non-substitutive designs (1.6 in Ebel *et al*., 2017 and 1.79 in Cryan *et al*., 2025). Net biodiversity effects were dominated by complementary effects, although selection effects still contributed to overall productivity. These findings are consistent with previous work showing that niche partitioning can promote overyielding in milpa systems (Postma & Lynch, 2012; Zhang et al., 2014). Importantly, the productivity advantage observed in the milpa was accompanied by changes in arthropod communities and by effects that persisted into the following growing season, suggesting that multiple processes may contribute to the benefits of crop diversification. Together, these results show the milpa to be a highly productive polyculture under low-input conditions where biodiversity can be particularly high (del-Val et al. 2021) .

Patterns in herbivore abundance and leaf damage did not consistently match productivity responses, suggesting that trophic interactions alone were not the primary drivers of productivity in this system, as it was shown in diversified cabbage systems (Croijmans et al. 2025). Instead, our findings indicate that complementary effects were more likely driven by plant-plant interactions, in particular belowground interactions related to resource acquisition and complementary resource use. The stronger productivity responses observed in squash and maize may reflect contrasting rooting strategies and differential soil resource exploitation among crops (Zhang et al. 2014). Nonetheless, ecological interactions are still likely to contribute to productivity under different environmental conditions or herbivore pressures. Previous work in the milpa system has shown that crop mixtures can enhance associational resistance and strengthen indirect defences against herbivores (Grof-Tisza et al. 2025), suggesting that inter-trophic ecological mechanisms may become particularly important when herbivore pressures are high. For example, bean extrafloral nectar can improve the attraction and fitness of parasitic wasps, potentially enhancing biological control across the three crops (Grof-Tisza et al. 2025). Likewise, squash shading and bean extrafloral nectar may provide additional ecological functions that become enhanced in polycultures.

Two factors may have contributed to the strong productivity differences observed in this study. First, the experiment was conducted under low fertilization, with nutrients added only twice during the season compared to five fertilizer applications typically used by local farmers. Since maize plants in the region require fertilizer to develop properly at the used densities, we decided to apply it moderately. This regime was intended to preserve potential contributions of nitrogen fixation by rhizobia symbiosis with beans and avoid homogenizing nutrient availability among plots. Under these conditions, competition for soil resources may have been stronger in monocultures than in polycultures. Second, bean plants developed poorly and produced very few pods, most likely due to intense herbivory by *Diabrotica* spp. This reduced bean performance may have indirectly favoured maize and squash growth in milpa plots by reducing competition among neighbouring plants. This may also explain why selection effects contributed relatively stronger to overall productivity compared to what has been previously estimated (Zhang et al. 2014). Even if low soil nutritional quality and differential plant-plant competition influenced the magnitude of the observed differences, the results clearly demonstrate that crop mixing can strongly enhance productivity under low-input conditions.

Herbivore abundance varied among crop species and cultivation types. Herbivore abundance was similar between monocultures and milpa plots on squash, higher in milpa on maize and higher in monoculture on beans. However, these patterns did not consistently match the intensity of leaf damage. Maize and squash experienced similar damage across diversity treatments, whereas bean plants suffered greater damage in milpa plots. This discrepancy likely reflects the poor development of beans during the experiment. Smaller bean plants supported fewer herbivores but suffered proportionally greater leaf damage. These results highlight the importance of indirect physical interactions among neighbouring plants in polycultures (Homulle et al. 2022). They also suggest that the yield outcome of crop mixing may depend strongly on crop species and genotype, environmental conditions, and plant variety, particularly when the species experience large differences in biotic stress (Li et al. 2020b; Schmutz and Schöb 2024). Predator abundance broadly mirrored herbivore abundance, suggesting that predator presence tracked prey availability, a common pattern in terrestrial food webs (Schmitz 2008). Importantly, the highest predator abundance was observed on maize plants in milpa plots. Maize may therefore act as an important focal crop for attracting natural enemies such as predatory social wasps which were frequently observed during the experiment. These wasps are known to respond to herbivore-induced maize volatiles (Southon et al. 2019; Grof-Tisza et al. 2024b), raising the possibility that maize in milpa systems could indirectly enhance biological control in neighbouring crops.

Herbivore diversity showed similar patterns to abundance which indicates that dominant species distributed evenly across the field. In contrast predator diversity was not influenced by our experimental factors. In a previous work it has been reported that milpa harboured a higher arthropod diversity than monocultures (Gorf-Tisza *et al.,* 2024a), which contrasts with the finding in this study that diversity was not consistently higher in milpa plots for all crop species. This can be explained by the fact that the sampling had been done previously at the plot level which means the three crop species contributed to arthropod diversity. In the present study diversity was estimated at the plant level thus it is not influenced directly by the three crop species. This indicates that the arthropod community in each crop species is affected differentially by increases in plant diversity, with maize community showing a positive response to milpa, whereas the opposite was observed for beans. In addition, seasonal variability could also account for changes in community responses to crop diversity (Nicholson 1958).

Beyond immediate productivity effects, our results suggest that crop diversity can influence the performance of subsequent crops through carryover effects that are consistent with soil legacy mechanisms (Jalloh et al. 2023; Ristok et al. 2023; Paranjape et al. 2026). Beans and squash both increased maize cob production the following season relative to sequential maize monoculture, while milpa plots showed intermediate responses. Although differences in final maize yield were not statistically significant, maize yield increased by nearly 30% in plots previously planted with beans and squash compared to continuous maize planting. This level of improvement can be quite relevant under agricultural conditions especially for smallholder farmers, for whom agronomic and economic benefits can be critical (Ricciardi et al. 2018). The results highlight the potential agronomic relevance of crop diversification and suggest that polycultures may complement crop rotation by generating benefits that persist into subsequent growing seasons. Crop rotation effects on soil fertility have been well documented (Zhao et al. 2020), and similar effects have been observed in other intercropping systems involving legumes (Xie et al. 2022; Jalloh et al. 2023). Maize can also benefit from microbial interactions mediated by secondary metabolites of neighbouring legume plants (Jiang et al. 2024). In addition, maize can produce negative effects on the yield of the next crop generation due to root exudates including benzoxazinoids and the associated microbiota (Cadot et al. 2021). Thus, heterospecific neighbours can help to prevent the build-up of these negative factors from maize. To our knowledge, however, this study represents one of the first field-based demonstrations that the milpa may influence subsequent crop performance through belowground carry over effects. These patterns are also consistent with parallel microbiome analyses conducted in the same experimental system, which suggest that crop diversity can alter belowground microbial communities associated with milpa plots (Rivera et al., submitted). Together, these findings suggest that plant-soil interactions may contribute to the carry over effects of previous crops observed in this study.

We initially expected beans to produce the strongest carry over effects because of their symbiosis with rhizobia and potential contribution to soil nitrogen availability. Interestingly, beans and squash produced similar responses. One possible explanation is that poor bean development limited root growth and nitrogen fixation during the experiment. In addition, a single growing season may not be enough for substantial nutrient accumulation and mineralization in the soil. Nitrogen transfer among crops may also be facilitated by long-term interactions among roots and mycorrhizal networks (Frankow-Lindberg and Dahlin 2013). However, soil nutrients, including nitrogen were not assessed in this study.

Taken together, our findings demonstrate that the milpa system can substantially increase productivity while influencing ecological interactions and generating benefits that extend beyond a single growing season (summarized in Figure 5). These findings are particularly relevant for smallholder farmers, who often have limited access to synthetic fertilizers, pesticides, and irrigation (Lowder et al. 2016). In these contexts, polycultures such as the milpa may provide meaningful agronomic benefits while simultaneously supporting biodiversity and maintaining traditional agricultural practices (Brooker et al. 2015; Novotny et al. 2021; Benrey et al. 2024). Maintaining natural habitats and arthropod diversity may be especially important in these systems because ecological interactions associated with biodiversity can contribute to pest regulation and crop performance (Liao et al. 2024; Grof-Tisza et al. 2025).

**Figure 5.**
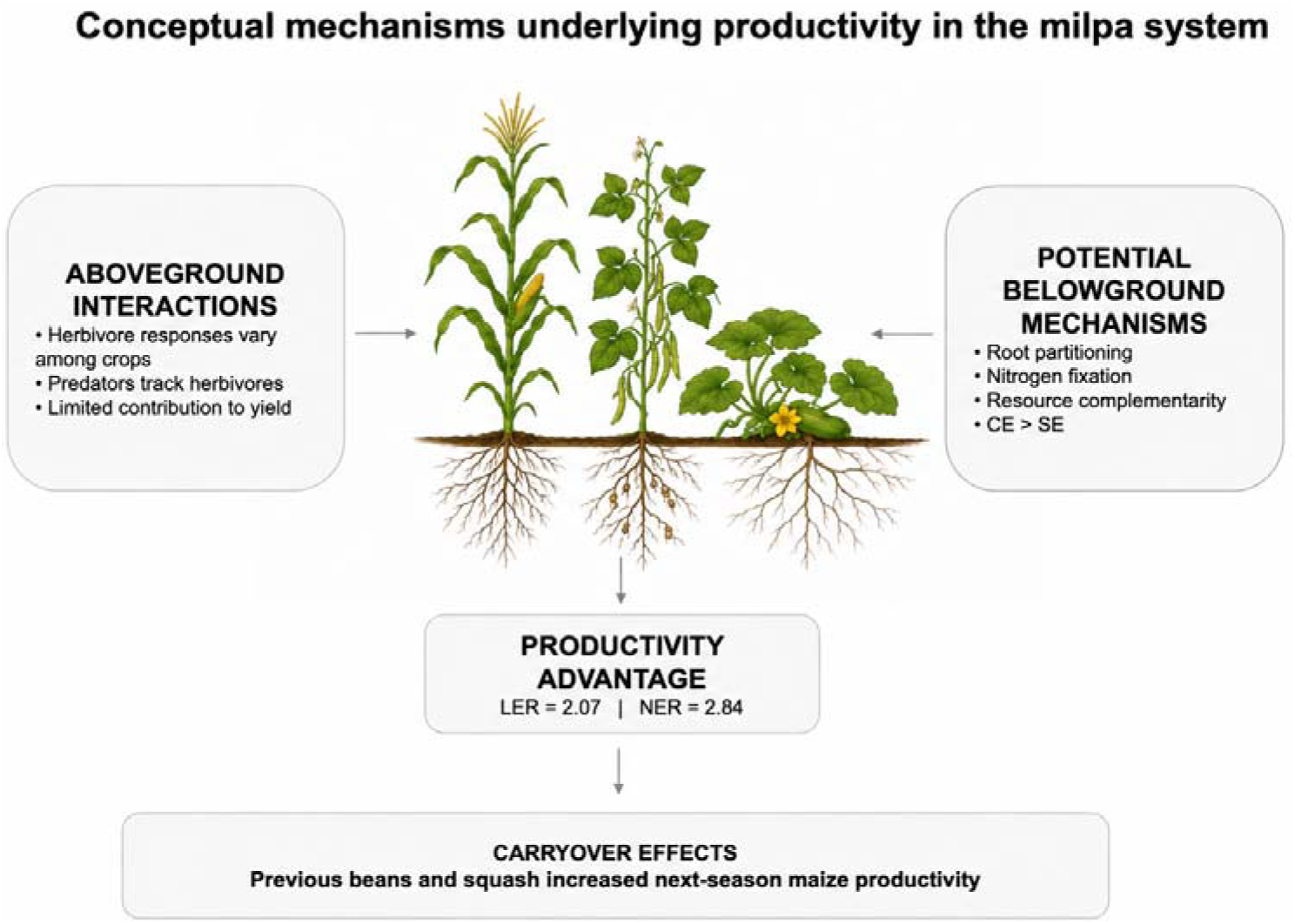
Conceptual summary of the main findings of this study. Milpa increased productivity primarily through complementarity effects. Although crop diversity altered arthropod communities, these changes did not explain productivity patterns under the conditions tested. Responses in the following growing season suggest potential carryover effects associated with previous crop composition. Proposed belowground mechanisms represent interpretations based on this study and previous work.

## Conclusions

Most studies on the milpa and other intercropping systems focus primarily on either productivity or biodiversity. In our study we examined how arthropod communities, herbivore damage, crop productivity, and responses in the subsequent growing season are linked under field conditions. The results suggest that the productivity advantage of the milpa is driven by complementary among crops rather than by changes in herbivore and predator abundance alone. This indicates that interactions among crops, including belowground resource partitioning, play an important role in generating productivity benefits in milpa polycultures. At the same time, ecological mechanisms such as associational resistance and biological control may contribute to crop performance under particular environmental conditions and should be investigated further. Finally, our findings suggest that the benefits of crop diversification may extend beyond a single growing season, emphasizing the importance of considering both immediate and longer-term consequences of crop mixing in sustainable low-input agriculture.

## DECLARATIONS

### Availability of data and material

The datasets and code generated during the current study are available in the Zenodo repository (https://doi.org/10.5281/zenodo.21772423)

### Authors’ contributions

Conceptualization, CBS, PGT and BB; Methodology, CBS, PGT, TCJT and BB; Investigation, CBS, PGT, CR, KG, RGS; Data analyses, CBS; Writing – Original Draft, CBS, PGT, BB; Writing –Review & Editing, All authors; Funding Acquisition, BB; Resources, BB; Supervision, CBS, PGT and BB

## Acknowledgements

We thank Clémence Nicollerat, Gaston Nobel, Loan Calame-Longjean and Sébastien Loup, for all their assistance in the field experiments. We are thankful to Alfredo Rojas for providing expert and logistic help in the field and the Rojas family for providing the field site.

